# Rewiring of EGFR oncogenic program by opposing actions of membrane versus soluble CD109 in HNSCC

**DOI:** 10.64898/2026.06.20.733552

**Authors:** Varsha R. Durgempudi, Tenzin Kungyal, Amani Hassan, Valentin Nelea, Kenneth Finnson, Dieter P. Reinhardt, Nader Sadeghi, Anie Philip

## Abstract

The epidermal growth factor receptor (EGFR) expression is often dysregulated in head and neck squamous cell carcinoma (HNSCC), driving cancer cell proliferation, invasion, and metastasis through diverse pathways, thereby contributing to aggressive chemo- and radio-therapy resistance. A GPI-anchored protein, CD109 is upregulated in multiple cancers, including HNSCC. While membrane-anchored CD109 (mCD109) is pro-tumorigenic in SCC via EGFR/STAT3 activation, the role of protease-cleaved soluble CD109 (sCD109) is poorly understood. Our groundbreaking findings demonstrate that sCD109 antagonizes EGFR signaling by directly binding to the EGFR extracellular domain, preventing mCD109-EGFR stabilizing interactions on the cell surface, followed by inhibition of EGFR phosphorylation at Y1068 and downstream signaling cascades (AKT, MAPK, and STAT3) consequently suppressing cancer cell migration, invasion, 3D tumor spheroid formation and angiogenic tube formation. In addition, we found that sCD109 regulates EGFR fates by inhibiting nuclear localization of phosphorylated EGFR and promoting EGFR degradation. Additionally, sCD109 significantly reduces EGF-induced expression of cancer stem cell markers (CD44 and CD133) and embryonic stem cell markers (Nanog and Sox2), suggesting a suppressive role in cancer stemness. Taken together, these results underscore the opposing roles of mCD109 and sCD109: with sCD109 acting as an antagonist by inhibiting mCD109/EGFR-driven oncogenic signaling and phenotypes. Our current findings reveal a complex interplay among mCD109, sCD109, and EGFR, identifying a mechanism for targeting EGFR’s degradation in HNSCC, and lay the groundwork for future research on investigating sCD109’s modulatory role in preclinical models of HNSCC.

## Introduction

Squamous cell carcinoma (SCC) originates from squamous epithelial cells in the outer layers of organs such as the skin, oral cavity, lungs, and cervix. Head and neck squamous cell carcinomas (HNSCCs) originate from the mucosal epithelium of the oral cavity, larynx, and pharynx (Sun et al., 2025). Currently, HNSCC is associated with a global incidence of more than 900,000 cases and a mortality of over 450,000 deaths every year (Xu et al., 2020). By 2040, GLOBOCAN estimates that HNSCC incidence will reach one million new cases and number of deaths are estimated to increase to 547,000 annually, making it the sixth most common cancer worldwide (Cancer, 2024). Despite advances in treatment modalities, HNSCC remains an aggressive malignancy, with 5-year survival declining from 88.4% in localized disease to 36.9% in distant metastatic disease (Institute, 2025). Its progression is characterized by increased proliferation, invasion, migration, epithelial-to-mesenchymal transition, and metastasis. While early-stage tumors may be effectively treated with surgery and radiotherapy, recurrence and distant metastasis are frequent, and effective strategies to curb these processes remain inadequate.

Aberrant activation of numerous oncogenic signaling pathways underlies the pathogenesis of HNSCC, with the EGFR axis occupying a central role. Binding of ligands such as EGF to the extracellular domain, induces receptor dimerization and autophosphorylation, and initiating downstream signaling cascades including the PI3K/Akt, Ras/MAPK, and STAT pathways that collectively promote proliferation, survival, migration, invasion, and metastasis. EGFR is overexpressed in the vast majority (approximately 90%) of HNSCCs and is closely linked to aggressive clinicopathological features, therapeutic resistance, and poor prognosis. In contrast, activating EGFR mutations are detected only in a subset of 10% of cases (Wang et al., 2009; Zhou et al., 2022). Although EGFR-targeted monoclonal antibodies such as cetuximab confer clinical benefit in selected patients, overall responses are modest and often short-lived because of intrinsic and acquired resistance. These limitations highlight an urgent need to delineate the molecular regulators of EGFR signaling and to develop more effective EGFR-directed inhibitors for HNSCC.

Cluster of differentiation 109 (CD109) is a membrane-anchored glycoprotein with broad tissue distribution. Our group was the first to identify a functional role for CD109, initially characterizing it as a TGF-β co-receptor (Bizet et al., 2011; Bizet et al., 2012; Finnson et al., 2006; Litvinov et al., 2011; Tam et al., 2001). CD109 belongs to the alpha-2-macroglobulin/C3/C4/C5 family and is expressed on platelets, T cells, endothelial cells, and keratinocytes (Lin et al., 2002; Man et al., 2012; Solomon et al., 2004; Vorstenbosch, Al-Ajmi, et al., 2013; Vorstenbosch, Gallant-Behm, et al., 2013; Vorstenbosch et al., 2017). Interestingly, CD109 is markedly upregulated in multiple malignancies, including squamous cell carcinomas (SCCs) of the head and neck (Tsutsumi et al., 2022; Zhou et al., 2017; Zhou et al., 2019; Zhou et al., 2022), lung (Hassan et al., 2025b; Lee et al., 2020; Taki et al., 2020), esophagus (Dong et al., 2015), urothelial tract (Hagikura et al., 2010), penis (Dong et al., 2017), cervix (Mo et al., 2020), as well as in glioblastoma (Filppu et al., 2021; Zhang et al., 2015), osteosarcoma (Mori et al., 2023), acute myeloid leukemia (Shyl et al., 2022), pancreatic (Haun et al., 2015), ovarian (Kim et al., 2019), breast (Hockla et al., 2010; Tao et al., 2014), and hepatocellular cancers (Cui et al., 2025; Ye et al., 2016). Recent studies by our group and others demonstrated that CD109 may promote tumor progression by enhancing the EGFR signaling pathway (Mo et al., 2020; Zhou et al., 2022) in SCC. More recently, we found that that CD109 is an important inhibitor of EGFR degradation in HNSCC and that EGFR membrane levels are critically dependent on CD109’s expression (Kungyal et al, 2026/BioRxiv).

We have previously reported that, like other GPI-anchored proteins, the CD109 ectodomain can be proteolytically shed from the cell surface by enzymes such as phosphatidylinositol-specific phospholipase C (PI-PLC) (Litvinov et al., 2011; Tam et al., 1998) to generate soluble CD109 (sCD109), which in turn can sequester the ligand TGF-β and antagonize TGF-β-induced Smad2/3 phosphorylation and fibrotic responses (Li et al., 2016; Tam et al., 2001). However, despite the established pro-tumorigenic role of mCD109, the impact of sCD109 on EGFR signaling and HNSCC progression has remained elusive. In the current study, we identify sCD109 as a mechanistic antagonist of EGFR signaling that counteracts mCD109-EGFR-driven oncogenic responses in HNSCC, thereby uncovering a previously unrecognized regulatory axis in EGFR-dependent cancer progression.

## Materials and methods

### Reagents

For signaling experiments, cells were serum-starved overnight and then left untreated or treated with 10 ng/ml EGF and/or 10nM sCD109 (Cat # 4385-CD-050 from R&D systems; Val22-Ser1268 with a C-terminal His-tag) for 15 minutes to measure phospho-EGFR (Y1068). To determine the effect of EGF on stemness, proliferation, and migration and 3D tumor spheroid formation, cells were left untreated or treated with 50 ng/ml EGF and/or 20nM sCD109 for 48 h. To determine the effect of EGF on 3D tumor spheroid formation, spheroids were left untreated or treated with 50 ng/ml EGF and/or 20nM sCD109 for 9 days, with replenishment every 3 days. For co-immunoprecipitation (Co-IP), cells were left untreated or treated with 10 ng/ml EGF and/or 10nM sCD109 for 15 minutes. Protein synthesis was inhibited by treating the cells with 0.5 mg/ml cycloheximide for different time points – 0, 1, 2, and 4 hours.

### Cell culture

The human head and neck squamous cell carcinoma cell lines - FaDu (derived from a hypopharyngeal SCC tumor of a 56-year-old male) and UMSCC38 (derived from a tonsillar SCC tumor of a 60-year-old male) cell lines were maintained in MEM and DMEM (with 1X MEM non-essential amino acids) supplemented with 10% fetal bovine serum respectively. Patient-derived primary cells were extracted from patient tissues as per the protocol.

### Collection of tumor tissue samples from HNSCC patients

HNSCC samples were surgically removed by Dr N. Sadeghi and his team at the Department of Otorhinolaryngology Head and Neck-Surgery, McGill University Health Centre. This study was approved by the Research Ethics Committee at the McGill University Health Centre (Protocol # MP-37-2019-4659) and conducted in accordance with the approved guidelines and regulations. All experimental protocols were approved by Research Review Office (RRO, Canada). Patients were advised of the procedures and provided written informed consent.

The eligibility criteria included previously untreated patients, HPV negative, without a second primary tumor and seeking treatment at the same institution. Stringent inclusion and exclusion criteria were applied and few of the inclusion criteria required a confirmed diagnosis of head and neck squamous cell carcinoma, age between 18 to 75, and no prior cancer therapy. Exclusion criteria included patients having nodal metastasis, severe comorbidities and/or pregnant/breastfeeding women. This study considered sex as a biological variable, ensuring an equitable representation of both female and male patients to attribute cancer progression and treatment outcomes differences. Blinding was performed to minimize bias, where possible, specifically in outcome assessments. Patient-derived primary cells were extracted from patient tissues as per our recent publication (Hassan et al., 2025a). Clinical data of patients is included in Supplementary Table 1.

### Surface plasmon resonance analysis

The SPR spectroscopy was performed to study the real-time interactions between sCD109-EGFR and sCD109-sCD109 self-interactions. The Biacore X instrument was used and sCD109 was covalently immobilised onto one of the two channels of the sensor chip by amine coupling. Binding experiments were performed by injecting 30 µL of analyte (EGFR/sCD109) at multiple concentrations. The dissociation constants K*_D_* were calculated as the average of individual K*_D_* values measured for each of the analyte concentrations tested. The experiments were repeated 2 times.

### Immunofluorescence

FaDu cells were plated onto 8-well chamber slides, were serum starved overnight and treated with 10 ng/ml EGF and/or 10 nM sCD109 for 15 minutes. Cells were washed with ice-cold PBS and were fixed with 4% paraformaldehyde for 20 mins at RT. Upon washing with PBS, cells were blocked with 4% bovine serum albumin in PBS for 30 mins and incubated with primary antibody overnight at 4°C. After PBS wash, cells were incubated with AF488 anti-mouse secondary antibody or AF594 anti-rabbit secondary antibody for 1 hour in the dark. Cells were washed with PBS and were mounted with coverslips of 1.4 thickness using mounting media, which includes Dapi. Samples were imaged using Zeiss LSM780 Laser Scanning Confocal Microscope at 63× (oil) objective. The experiments were repeated 3 times.

### Proximity ligation assay

Duolink In Situ Red Starter Kit Mouse/Rabbit from Sigma was used to measure the PPIs between mCD109-EGFR, sCD109-EGFR, and sCD109-mCD109. The protocol followed explicitly the manufacturer’s instructions, and 8-well chamber slides were used for primary cells. Samples were imaged using Zeiss LSM780 Laser Scanning Confocal Microscope at 63× (oil) objective. The experiments were repeated 3 times.

### Co-immunoprecipitation

SureBeads (Bio-Rad) or Protein G magnetic beads (Genscript) were thoroughly resuspended, and 100 µl aliquots were transferred to 1.5ml Eppendorf tubes. The supernatant was discarded upon magnetization. Beads were washed with PBST (0.1% Tween20) three times. About 2 µg of anti-EGFR antibody in a final volume of 200 µl was added to the beads, resuspended, and rotated on an end-over-end shaker for 1 hour at RT. The supernatant was discarded upon magnetization. Beads were washed with PBST (0.1% Tween20) three times. Cells were left untreated or treated with 10 ng/ml EGF and/or 10nM sCD109 for 15 minutes and harvested in IP-lysis buffer. The antigen-containing lysate (about 400 - 600 µg) was added to the beads and rotated on an end-over-end shaker overnight at 4°C. The supernatant was discarded upon magnetization. Beads were washed with PBST (0.1% Tween20) three times. The supernatant was discarded upon magnetization after transferring the resuspended beads to a new tube. About 40 µl of SDS sample buffer was added to the beads and incubated at 70°C for 10 min. Beads were magnetized, and the eluent was subjected to western blotting to probe with anti-CD109 antibody. Reciprocal co-IP experiments were conducted as well. The experiments were repeated 3 times.

### Western blot analysis

FaDu cell lysates (containing 40 µg total protein) were analyzed by western blotting with one of the following antibodies: mouse anti-CD109, anti-EGFR, anti-phospho-STAT3, anti-STAT3, anti-phospho-AKT, anti-AKT, anti-phospho-ERK, anti-ERK, anti-Nanog, anti-CD44, anti-SOX2, rabbit anti-phospho-EGFR, and anti-β-actin antibodies. Briefly, western blot analysis involved cell lysates harvesting in RIPA buffer, incubated at 95°C for 10 min with SDS sample buffer, followed by running lysates by SDS polyacrylamide gel electrophoresis and transferring them to nitrocellulose membranes. These membranes were washed with TBST (0.1% Tween20) buffer and stained with Ponceau S solution for 10 min at room temperature (RT). Then images were acquired and washed with TBST buffer to destain. They were blocked with 1% bovine serum albumin/TBST buffer at RT for 45 min and then incubated overnight with respective primary antibodies at 4°C. The blots were washed with TBST buffer and incubated with appropriate HRP-conjugated secondary antibodies for 1 hour at RT. The blots were developed with an enhanced chemiluminescence ECL system.

### Flow cytometry

FaDu cells were left untreated or treated with 10 ng/ml EGF and/or 10nM sCD109 for 15 minutes, harvested by Accutase, washed with ice-cold PBS, and then resuspended in staining buffer (2% FBS, 1mM EDTA in PBS), blocked with blocking buffer (2% FBS in PBS), and were processed for Flow cytometry analysis using anti-EGFR tagged with AF-647 to check for membrane EGFR expression in cells. Sample were then analyzed using BD LSR Fortessa 4L. The experiments were repeated 3 times.

### Surface protein biotinylation

FaDu cells were left untreated or treated with 10 ng/ml EGF and/or 10nM sCD109 for 15 min. Media was aspirated from the plate, and cells were rapidly washed twice with 3 ml of ice-cold PBS in each well. The following steps were carried out on the ice. Two vials (1mg each) of Biotin were instantly dissolved in 8 ml of PBS to make a final 250 µg/ml concentration and were distributed by 1300 µl into each well. The plate was placed on an orbital shaker for 30 min at 4°C. About 65 µl of Quenching solution was added to the plate wells each and incubated for 5 min at 4°C after ensuring even exposure. Cells were gently scraped, and solutions were transferred into 1.5 ml Eppendorf tubes. Wells were rinsed with 640 µl of ice-cold TBS and were pooled into the same tubes. Cells were spun at 500×g for 3 min, and the supernatant was discarded. About 640 µl of TBS was added onto the cell pellet and gently resuspended. Cells were spun at 500×g for 3 min, and the supernatant was discarded. The TBS washing step was repeated once again. A protease inhibitor cocktail was added instantly to the Lysis solution, and about 90 µl of Lysis solution was added to the cell pellet. After proper resuspension, samples were incubated on an end-over-end shaker for 1 hour at 4°C. Medium speed vortexing was performed every 10 min for 5 s. Lysates were spun at 10,000×g for 15 min at 4°C. Supernatants (cell lysates) were collected into new tubes. Bradford protein estimation was done, and about 40 µg of protein were taken out as whole-cell lysate control.

About 80 µl of streptavidin agarose slurry was taken for each sample in multiple columns and were spun at 2,000×g for 1 min. The supernatant was discarded and washed with about 100 µl of washing buffer at 2,000×g for 1 min thrice. Before the third spin, slurries were transferred to Eppendorf tubes, and the supernatant was removed slowly using gel loading tips. Equal amounts of cell lysates were added onto slurries and incubated on an end-over-end shaker overnight at 4°C. Samples were short-spun, transferred to fresh columns, then spun at 2,000×g for 1 min. This supernatant was collected into new Eppendorf tubes and used as flow-through controls. A protease inhibitor cocktail was added instantly to the washing buffer, and about 100 µl of washing buffer was added to columns, followed by spinning at 2,000×g for 1 min three times. About 50 µl of PBS was used to resuspend the resins, and they were transferred to fresh Eppendorf tubes. Samples were spun at 2,000×g for 1 min, and the supernatant was removed slowly using gel loading tips. About 20 µl of 1M dithiotreitol was instantly added to 400 µl of RIPA buffer (diluted with sterile water), and about 60 µl of this Elution mix was added to the precipitated resins, followed by incubation on an end-over-end shaker at 37°C for 1 hour. Resins were spun at 2,000×g for 2 min, and supernatants were subjected to western blotting. The experiments were repeated 3 times.

### Immunocytochemistry

FaDu cells were left untreated or treated with 10 ng/ml EGF and/or 10 nM sCD109 for 15 min. Cells were washed with ice-cold PBS and were fixed with 4% paraformaldehyde for 20 min at ambient temperature. Upon washing with PBS, endogenous peroxidase blocking was done for 5 min at RT, followed by blocking with 4% bovine serum albumin in PBST (0.1% TritonX100) for 1 hour. Primary antibody incubations were overnight at 4°C with mouse anti-phospho-EGFR antibodies. Cells were washed with distilled water three times and then incubated with mouse complement for 10 min, followed by water wash and HRP-conjugated secondary antibody for 15 min. Reactions were developed with DAB chromogen and substrate, then counter-staining with Hematoxylin for 5 min. Cells were subjected to acid-alcohol wash using HCl and 95% ethanol for 8-10 dips. Cells were dehydrated with 100% ethanol and xylene, followed by mounting with Cytoseal mounting media. The experiments were repeated 3 times.

### Fluorescent western blotting

FaDu cell lysates were prepared like a chemiluminescent western blotting and gels were run using “low-fluorescence protein marker”. Wet transfers were done using “low fluorescence” PVDF membranes (after pre-wetting in methanol for 30 s, briefly rinsing with water, and equilibrating in transfer buffer for 5 min). Blots were washed with washing buffer for 10 min followed by blocking in 4% BSA in TBST (0.05% Tween 20) for 30 min. Primary antibody incubations were done overnight. Followed by 3 washes, secondary antibody incubations were done with one fluorophore for 1 hour at RT (all steps from now in dark). Followed by 3 washes, a second primary antibody incubations were performed for 1 hour at RT. Followed by 3 washes, secondary antibody incubations were done with another fluorophore for 1 hours at RT. Followed by 3 washes, fluorescence detection was performed. All buffers used were filter sterilized. The experiments were repeated 3 times.

### Nuclear extraction

FaDu cells were left untreated or treated with 10 ng/ml EGF and/or 10 nM sCD109 for 15 min. Subcellular protein fractionation kit protocol from ThermoFisher was explicitly used to fractionate cytosolic and nuclear fractions from HNSCC cells. Both fractions were further analyzed by western blotting to probe for phospho-EGFR, Actin and Lamin-B1. The experiments were repeated 3 times.

### Cycloheximide chase assay

FaDu cells were treated with Cycloheximide (0.5 mg/ml) and/or 10 nM sCD109 at 0, 1-, 2-, 4- and 8-hour time points and were further analyzed by Western blotting to probe for EGFR, CD109 and Actin. The experiments were repeated 3 times.

### MTS cell proliferation assay

FaDu cells were left untreated or treated with 50 ng/ml EGF and/or 10nM sCD109 for 24- and 48-hour time points, followed by incubation with MTS reagent for 4 hours at 37°C. The plate was briefly shaken on a shaker and the OD was measured at 490 nm for quantifying cell proliferation. The experiments were repeated 3 times.

### In vitro scratch (circular wound healing) assay

FaDu and UMSCC38 cells cultured in a 24-well plate were left untreated or treated with 50 ng/ml EGF and/or 10 nM sCD109 for 48 hours.

We modified the protocol from (De Ieso & Pei, 2018). A sterile 200µl tip was inserted into the vacuum aspirator collection tube and a circular scratch was made by touching the base of the plate. Several such wounds were made in a pattern and were labelled to further monitor the wound healing area after 24 and 48 hours of treatment. Images were captured using an EVOS inverted microscope at 10× and 20× objective and Fiji (ImageJ) software (Schindelin et al., 2012) was used to measure the width of the wound area. The experiments were repeated 3 times. Cell migration was expressed as percentage of the scratch area filled by migrating cells at 24 h/48 h post scratch: migration rate = (T0 h scratch width − T24/T48 h scratch width)/T0 h scratch width) × 100%.

### Matrigel cell invasion assay

Matrigel was diluted from stock (10 mg/ml) to 300 µg/ml and added about 100 µl on top of sterile 24-well cell culture inserts followed by incubation in pre-heated 24-well plates at 37°C for 2 hours. About 7×10^4^ FaDu cells in basal media were added on top of solidified Matrigel per insert. Cells were allowed to attach on solidified Matrigel at 37°C for 4 hours. After gently aspirating the media on top, cells were pre-incubated with EGFR inhibitors at 37°C for 45 min. EGF was added to bottom chambers in 24 wells as a chemoattractant in respective wells and were incubated at 37°C for 24 hours. After 24 hours, inserts were gently washed with PBS (both top and bottom sides), fixed with 4% PFA and were stained with Dapi (1ug/ml) for 10 min in dark at RT. Bottom side of inserts were then imaged using an EVOS inverted microscope at 4× and 10× objective. The experiments were repeated 3 times. The total number of Dapi-stained cells were counted using Image-J software.

### 3D Tumor spheroids assay

About 10,000 FaDu cells were cultured in an ultra-low attachment 6-well plate for 3 days in 3ml of DMEM/F12 media with growth factors and supplements (20 ng/ml recombinant human fibroblast growth factor-10 (FGF10), 20 ng/ml hFGF2, 9.5 µg/ml insulin and 2% Gibco B-27 Supplement). On day 3, 1ml of spheroid media was topped up along with treatments of 50 ng/ml EGF and/or 10 nM sCD109. Topping up with 1ml of spheroid media along with treatments were repeated on day 6 and day 9. Images were captured and individual spheres were counted using an EVOS inverted microscope at 10× and 20× objective. The experiments were repeated 3 times. The percentage of cells capable of forming spheres was calculated as follows: [(number of spheres formed/number of cells plated) × 100].

### Angiogenesis tube formation assay

About 100 µl of Matrigel was added to pre-heated 48-well cell culture plates and were incubated at 37°C for 1 hour. About 1.2×10^5^ HUVEC cells were cultured on top of solidified Matrigel in basal media along with/without EGF and sCD109. The plate was incubated at 37°C for 18 hours. After 18 hours, culture media was gently aspirated, washed with PBS and stained with 1mM CMFDA stain at 37°C for 10-15 min. The stain was removed and PBS was added to further image the tube formation using EVOS inverted microscope. The experiments were repeated 3 times. The total length of the tubes (in µm) were quantified using Image-J software.

### Statistical analysis

All quantitative data are presented as mean ± SD. Data was analyzed by two-tailed Student’s t-test or one-way and two -way Analysis of Variance (ANOVA). A p-value < 0.05 was considered significant.

## STAR★Methods

### Key resources table

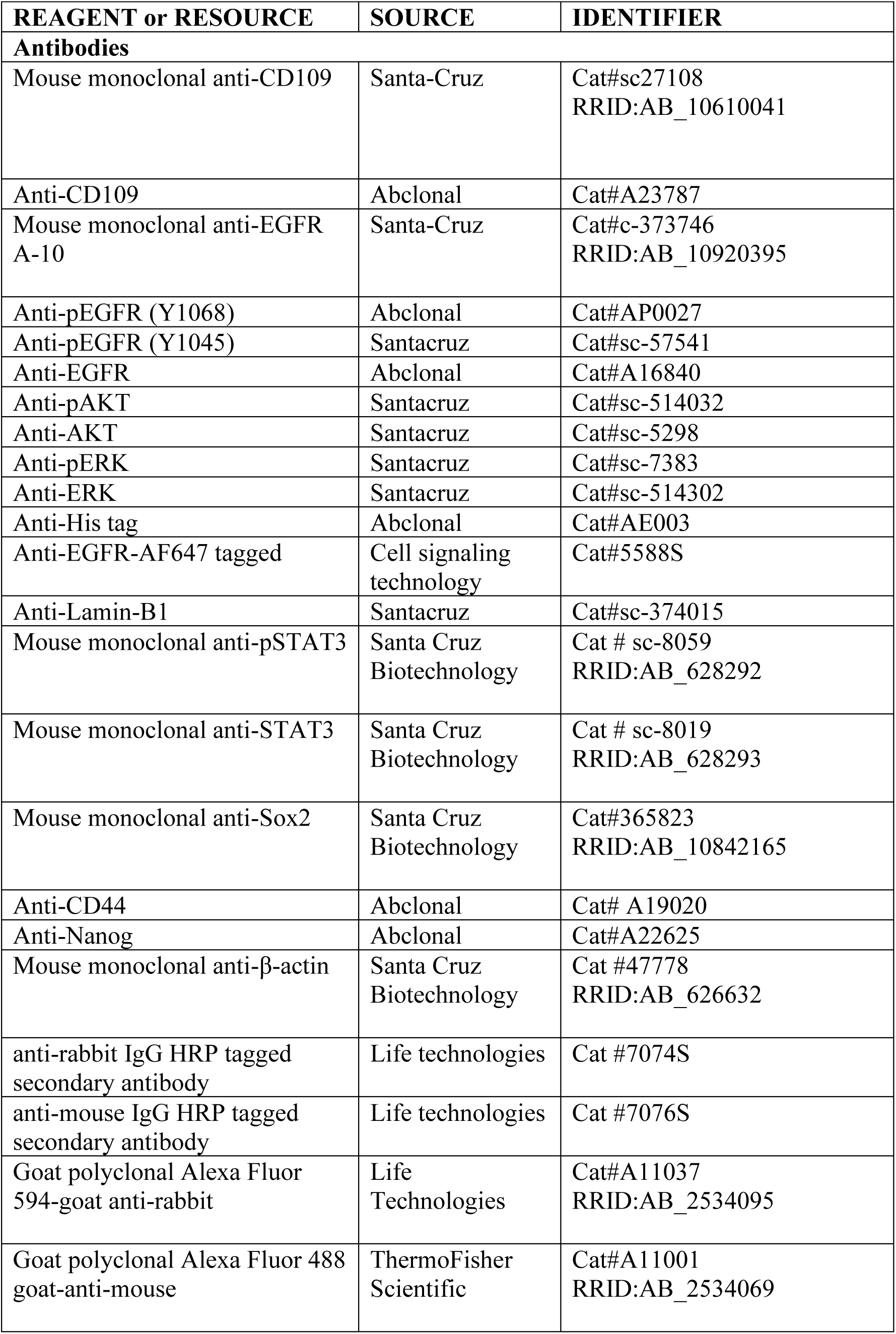
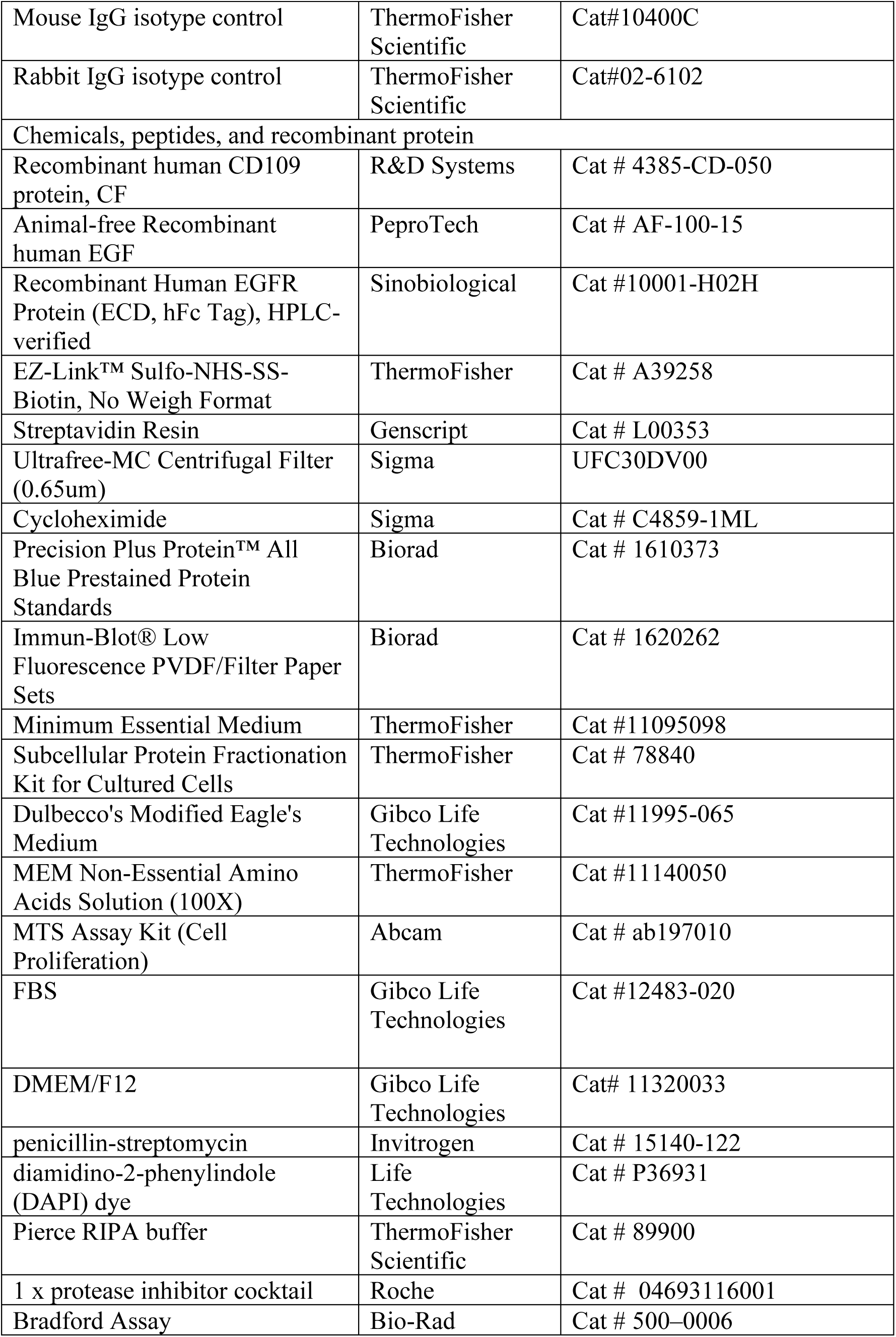
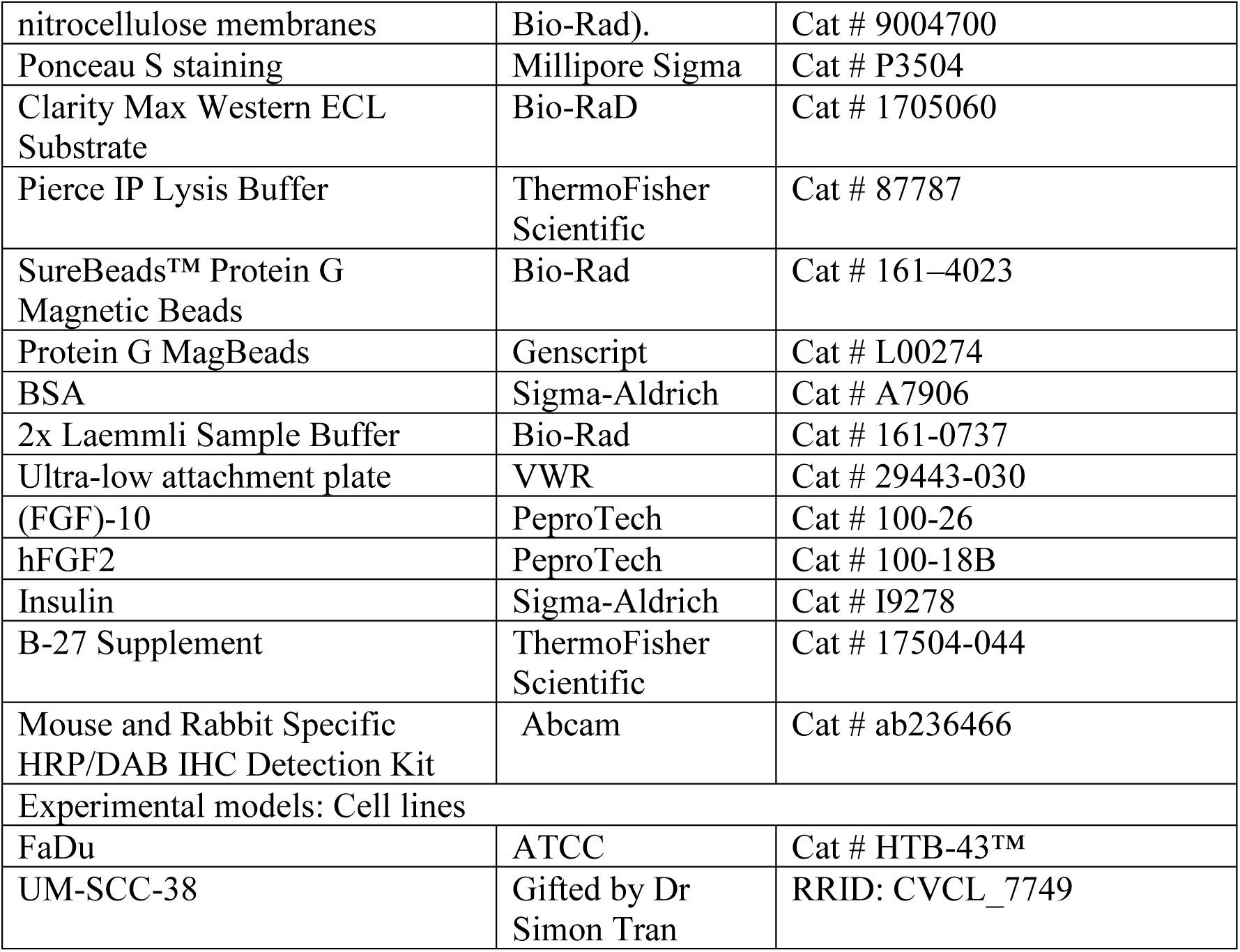

### Results

#### Membrane CD109 associates with EGFR and stabilizes EGFR cell surface levels while soluble CD109 competitively inhibits this association

EGF time-course experiments established 15 min as the peak window for detecting phosphorylation at Y1068 and Y1045 sites. (Supplementary Fig. 1A). To establish the role between mCD109 and EGFR in HNSCC cells, we showed that mCD109 overexpression promotes EGFR stabilization and phosphorylation in SCC9 mCD109 OE and EV cell lines. In addition, we showed that mCD109 KO led to decreased EGFR levels and inhibited phosphorylation in SCC9 KO cells when compared to WT cells (Fig. 1A). To delineate the role of sCD109 in regulating the mCD109-EGFR stabilizing interactions, we performed PLA in patient-derived primary cells. The PLA analysis demonstrated that sCD109 treatment markedly reduced EGF-induced mCD109-EGFR interactions (Fig. 1B). The co-immunoprecipitation analysis in FaDu cells confirmed this sCD109’s competitive inhibitory role on EGF-induced mCD109-EGFR complex (Fig. 1C). Preliminary binding assays confirmed direct association of sCD109 and EGFR (Supplementary Fig. 1B). To investigate the direct binding of sCD109 (contains Histag) to EGFR compared to its membrane counterpart, we performed PLA using anti-Histag antibody with anti-EGFR and anti-CD109 antibodies in patient-derived primary cells. We showed that sCD109’s interactions with EGFR were significantly higher than with mCD109, both in the presence and absence of EGF (Fig 1D). The SPR analysis exemplified that sCD109 binds the EGFR ectodomain with picomolar affinity (KD = 78 ± 15 pM), about 2000-fold stronger than CD109 self-interaction (KD = 155 ± 67 nM) (Fig. 1E). Immunofluorescence microscopy revealed significant co-localization of His-tagged sCD109 with EGFR at the cell surface using anti-Histag antibody and anti-EGFR antibody (Fig. 1F). Altogether, these data demonstrate that mCD109 stabilizes EGFR expression while sCD109 binds to EGFR ectodomain with high affinity and competitively displaces mCD109 from interacting with EGFR in HNSCC cells.

**Figure 1:**
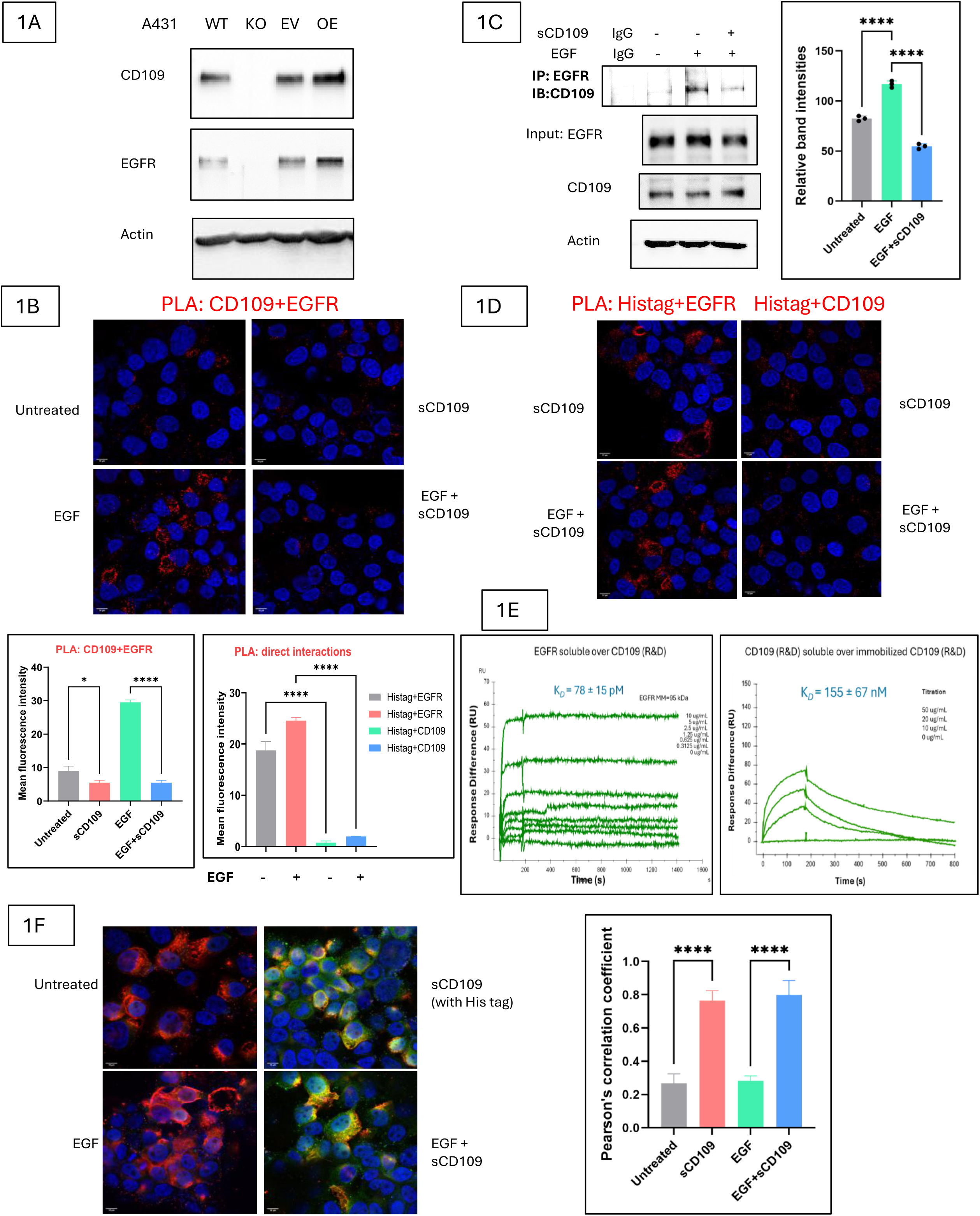
Membrane CD109 associates with EGFR and stabilizes EGFR cell surface levels while soluble CD109 competitively inhibits this association. A) Western blotting analysis of phospho-EGFR (Y1068), EGFR and CD109 upon treatment of SCC9 CD109 OE, EV, WT and KO cells with EGF for 15 mins. B) PLA to detect interactions between mCD109 and EGFR and the role of sCD109 treatment to these interactions in patient-derived primary cells. Cells were treated with EGF and/or sCD109 for 15 mins. C) Co-immunoprecipitation and western blotting analysis to check the role of sCD109 on the interactions of mCD109 and EGFR in FaDu cells. Cells were treated with EGF and/or sCD109 for 15 mins and mouse IgG control is also included. D) PLA to detect interactions between sCD109 and EGFR [left panel] and sCD109 and CD109 [right panel], both in the presence/absence of EGF in patient 1’s primary cells. Cells were treated with EGF and/or sCD109 for 15 mins. E) SPR spectroscopy analysis of sCD109 vs EGFR and itself (self interactions). F) Immunofluorescence co-localization studies of sCD109 (anti-His tag antibody - green) and EGFR (anti-EGFR antibody - red) both in the presence/absence of EGF in FaDu cells. Cells were treated with EGF and/or sCD109 for 15 mins. All the results were expressed as the mean+/- SD of three independent experiments, except B), D) and E) which were expressed as the mean+/- SD of two independent experiments. Significance was calculated using one-way ANOVA *p < 0.01.

#### Soluble CD109 reduces membrane-anchored and total EGFR levels in HNSCC cells

Based on our prior findings that mCD109 stabilizes EGFR at the plasma membrane, we next examined whether sCD109 alters receptor expression levels. Immunocytochemistry revealed a decrease in total EGFR levels upon sCD109 treatment for 15 mins of treatment time (Fig. 2A). Flow cytometry using anti-EGFR-PE antibody in FaDu cells detected a time-dependent reduction in surface EGFR levels (0–30 min) upon sCD109 treatment in basal conditions (Fig. 2B). To validate these findings, we performed cell-surface biotinylation of receptors for 30 mins followed by streptavidin pull-down and western blotting to probe for EGFR. We observed that sCD109 treatment for 15 mins reduced membrane-localized EGFR under both basal and EGF-induced conditions (Fig. 2C). Altogether, these data demonstrate that sCD109 treatment decreases both surface and total EGFR expression in FaDu cells, consistent with disruption of receptor stabilization and enhanced receptor downregulation.

**Figure 2:**
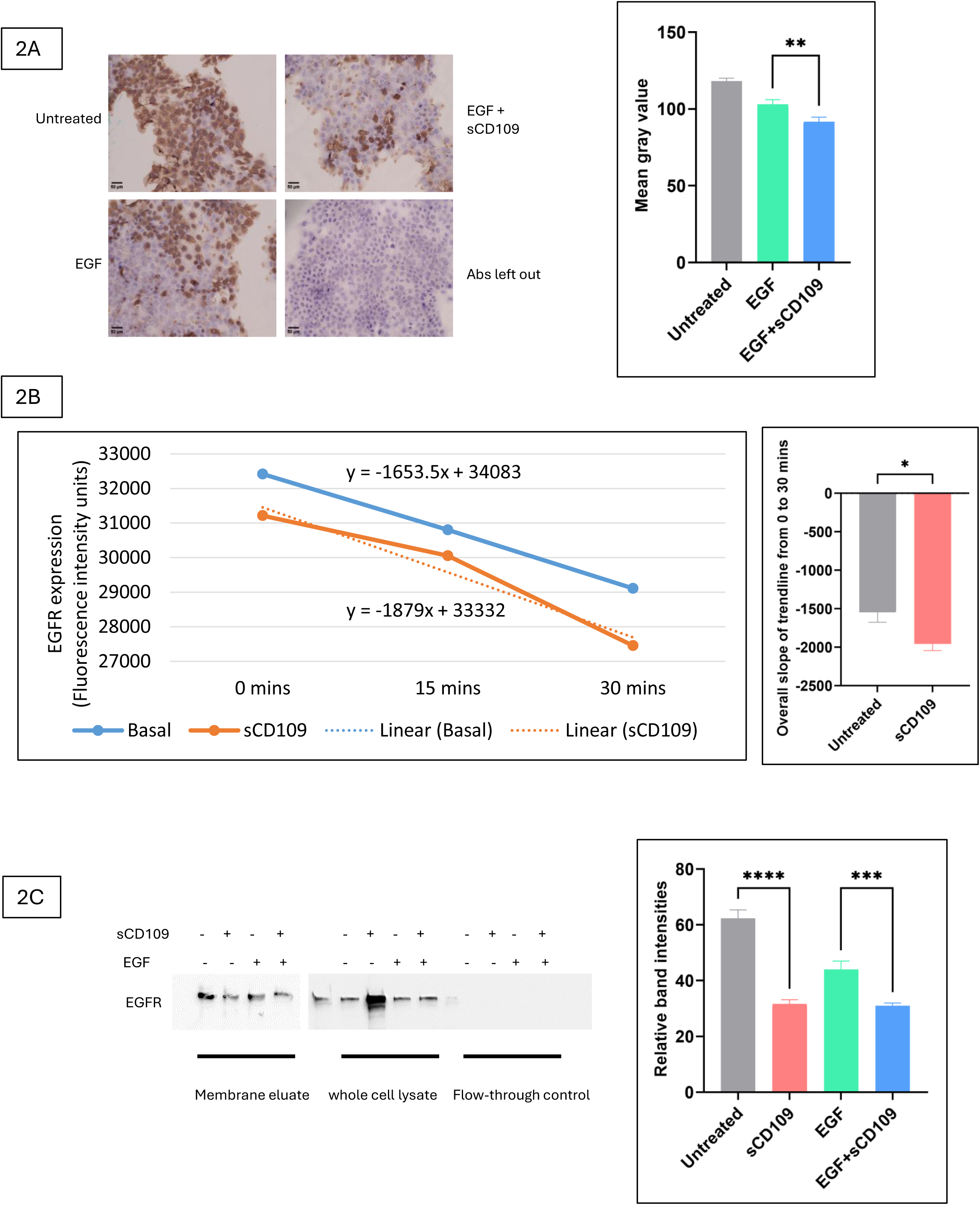
Soluble CD109 reduces membrane-anchored and total EGFR levels in HNSCC cells. A) Immunocytochemistry analysis of total EGFR expression upon treatment of FaDu cells with EGF and/or sCD109 for 15 mins. B) Flow cytometry analysis of cell surface EGFR upon treatment of FaDu cells with sCD109 at multiple time points of 0, 15 and 30 mins. C) Biotinylation and streptavidin pull-down and western blotting analysis of cell surface EGFR upon treatment of FaDu cells with EGF and/or sCD109 for 15 mins. All the results were expressed as the mean+/-SD of three independent experiments. Significance was calculated using one-way ANOVA *p < 0.01.

#### Soluble CD109 attenuates EGF-induced EGFR/STAT3, AKT, and ERK signaling by reducing signaling-associated EGFR Y1068 phosphorylation while enhancing degradation-associated Y1045 phosphorylation

We next investigated the role of sCD109 binding to EGFR on its activation. Dose-response analyses in A431, FaDu cells and patient-derived primary HNSCC cells identified 10 nM as the optimal concentration for phospho-EGFR and downstream signalling readouts (Supplementary Fig. 1C, D, E). Treatment of HNSCC cells with sCD109 significantly inhibited the EGF-induced phosphorylation of EGFR at Y1068 epitope upon 15 mins of treatment in FaDu and UMSCC38 cells (Fig. 3A; Supplementary Fig. 1F). The potency of sCD109 inhibition was comparable to established EGFR-inhibitors, including cetuximab and AG1478 (Supplementary Fig. 1G). Subsequent downstream signaling analyses showed sCD109’s inhibition of phospho-STAT3 at Y705, phospho-AKT at S473, and phospho-ERK at Y204 epitopes in FaDu cells and patient-derived primary cells (Fig. 3B,C). The immunocytochemistry and immunofluorescence experiments using anti-pEGFR and anti-EGFR antibodies (Fig. 3D,E), confirmed that sCD109 downregulates EGFR-mediated signaling axis activation in HNSCC progression.

**Figure 3:**
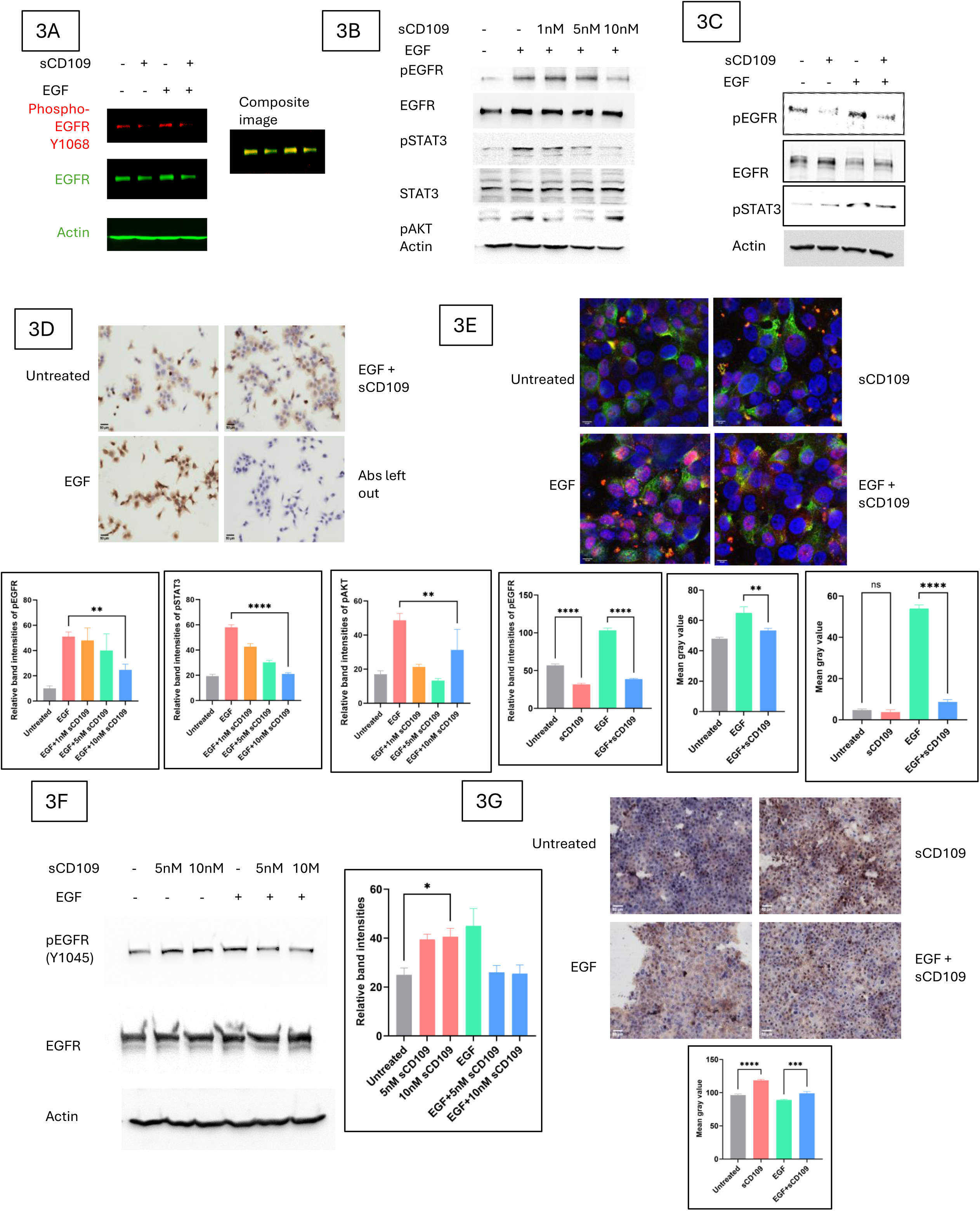
Soluble CD109 attenuates EGF-induced EGFR/STAT3, AKT, and ERK signaling by reducing signaling-associated EGFR Y1068 phosphorylation while enhancing degradation-associated Y1045 phosphorylation. A) Fluorescent western blotting analysis of phospho-EGFR (Y1068) (red) and total EGFR (green) upon treatment of FaDu cells with EGF and/or sCD109 for 15 mins. B) Western blotting analysis of phospho-EGFR (Y1068), phospho-STAT3 and phospho-AKT upon treatment of FaDu cells with EGF and/or sCD109 for 15 mins. C) Western blotting analysis of phospho-EGFR (Y1068) and phospho-STAT3 upon treatment of patient 1’s primary cells with EGF and/or sCD109 for 15 mins. D) Immunocytochemistry analysis of phospho-EGFR (Y1068) upon treatment of FaDu cells with EGF and/or sCD109 for 15 mins E) Immunofluorescence analysis of phospho-EGFR (Y1068) (red) and EGFR (green) upon treatment of FaDu cells with EGF and/or sCD109 for 15 mins. F) Western blotting analysis of phospho-EGFR (Y1045) upon treatment of FaDu cells with EGF and/or sCD109 for 15 mins. G) Immunocytochemistry analysis of phospho-EGFR (Y1045) upon treatment of FaDu cells with EGF and/or sCD109 for 15 mins. All the results were expressed as the mean+/- SD of three independent experiments. Significance was calculated using one-way ANOVA *p < 0.01.

In contrast to its role at Y1068, sCD109 promoted phosphorylation at Y1045 epitope upon 15 mins of treatment under basal conditions, a site linked to c-Cbl recruitment and EGFR ubiquitination, observed via western blotting and immunocytochemistry (Fig. 3F,G). Taken together, these findings show that sCD109 inhibits EGF-induced EGFR phosphorylation at Y1068 while promoting EGFR phosphorylation at Y1045 epitopes in FaDu cells.

#### Soluble CD109 impairs nuclear translocation of phospho-EGFR and accelerates EGFR degradation

To investigate the role of sCD109 on EGFR trafficking and fates, we investigated nuclear localization of phosphorylated EGFR which is often associated with poor prognosis in HNSCC. Nuclear fractionation of FaDu cells followed by immunoblotting with phospho-EGFR and LaminB1 as nuclear loading control revealed that sCD109 decreased EGF-induced nuclear translocation of phospho-EGFR (Fig. 4A), a finding confirmed by immunofluorescence imaging (Fig. 4B).

**Figure 4:**
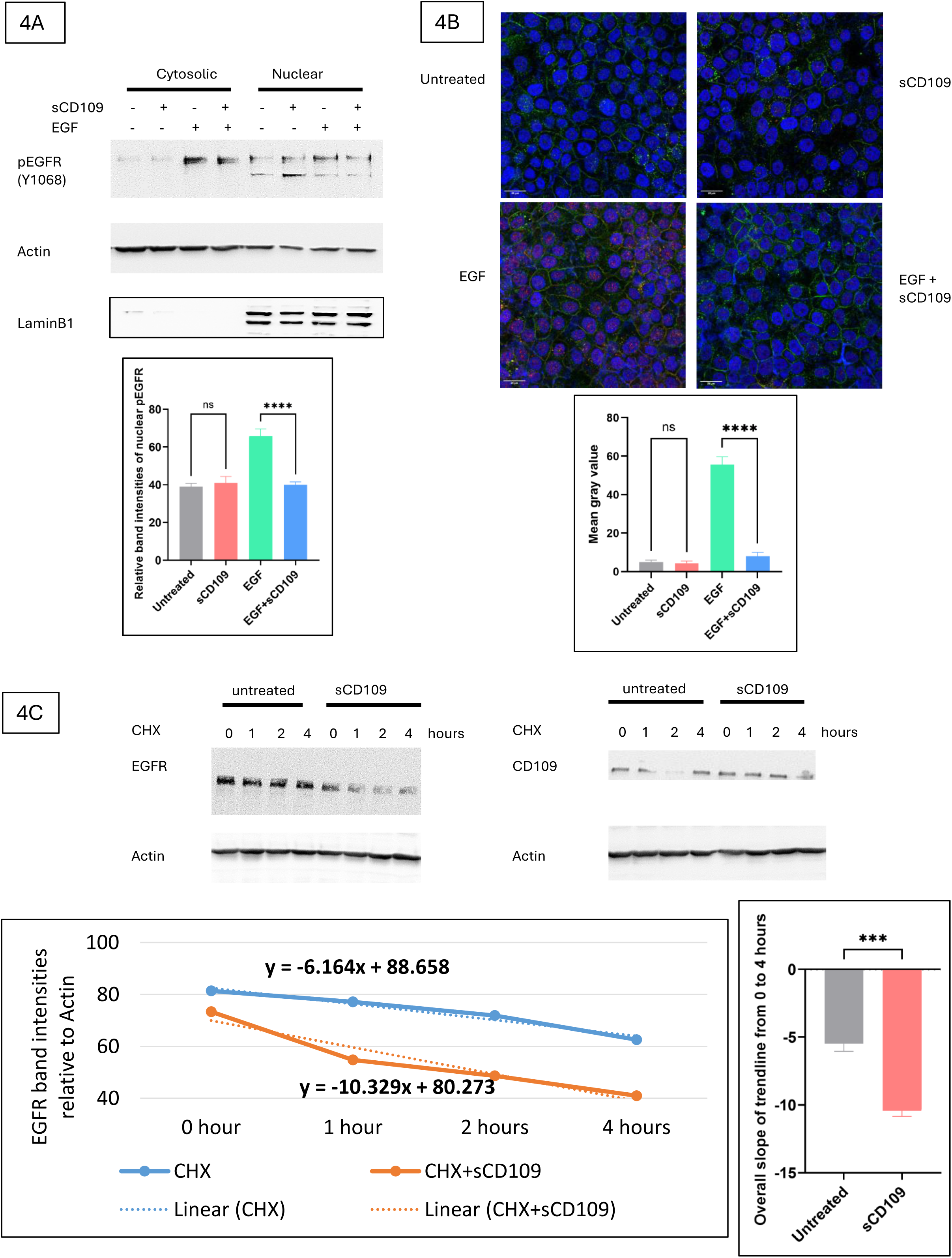
Soluble CD109 impairs nuclear translocation of phospho-EGFR and accelerates EGFR degradation. A) Nuclear extraction and western blotting analysis of phospho-EGFR (Y1068) upon treatment of FaDu cells with EGF and/or sCD109 for 15 mins. Actin was used as a cytosolic control and LaminB1 as a nuclear control. B) Immunofluorescence analysis of phospho-EGFR (Y1068) upon treatment of FaDu cells with EGF and/or sCD109 for 15 mins, followed by Tween20 permeabilization of cells. C) Cycloheximide chase (CHX) assay and western blotting analysis of EGFR upon treatment of FaDu cells with sCD109 for multiple time points of 0, 1, 2, and 4 hours. Quantification of overall slope was also performed. All the results were expressed as the mean+/- SD of three independent experiments. Significance was calculated using one-way ANOVA *p < 0.01.

A time-dependent cycloheximide chase revealed accelerated degradation of EGFR in sCD109-treated cells within 4 hours of treatment time under EGF-unstimulated conditions (Fig. 4C). Together, these results demonstrate that sCD109 restricts EGFR signaling not only by blocking activation at the membrane but also by preventing nuclear trafficking and promoting receptor degradation.

#### Soluble CD109 suppresses stemness-associated transcriptional programs in HNSCC

In addition to investigating the role of sCD109 on EGFR expression, signaling and cellular fates, we investigated the expression of widely known cancer stem cell markers and embryonic stem cell markers in HNSCC progression. Following 48 hours of treatment with sCD109 in FaDu cells, western blotting revealed significant downregulation of CD44, CD133, and SOX2 in basal and EGF-induced conditions (Fig. 5A) and CD44, Nanog and SOX2 in patient 2’s primary cells (Figure 5B). Overall, these findings demonstrate that sCD109 inhibits the EGF-induced expression of CD44, Nanog, and Sox2 in HNSCC cell lines and patient-derived primary cells.

**Figure 5:**
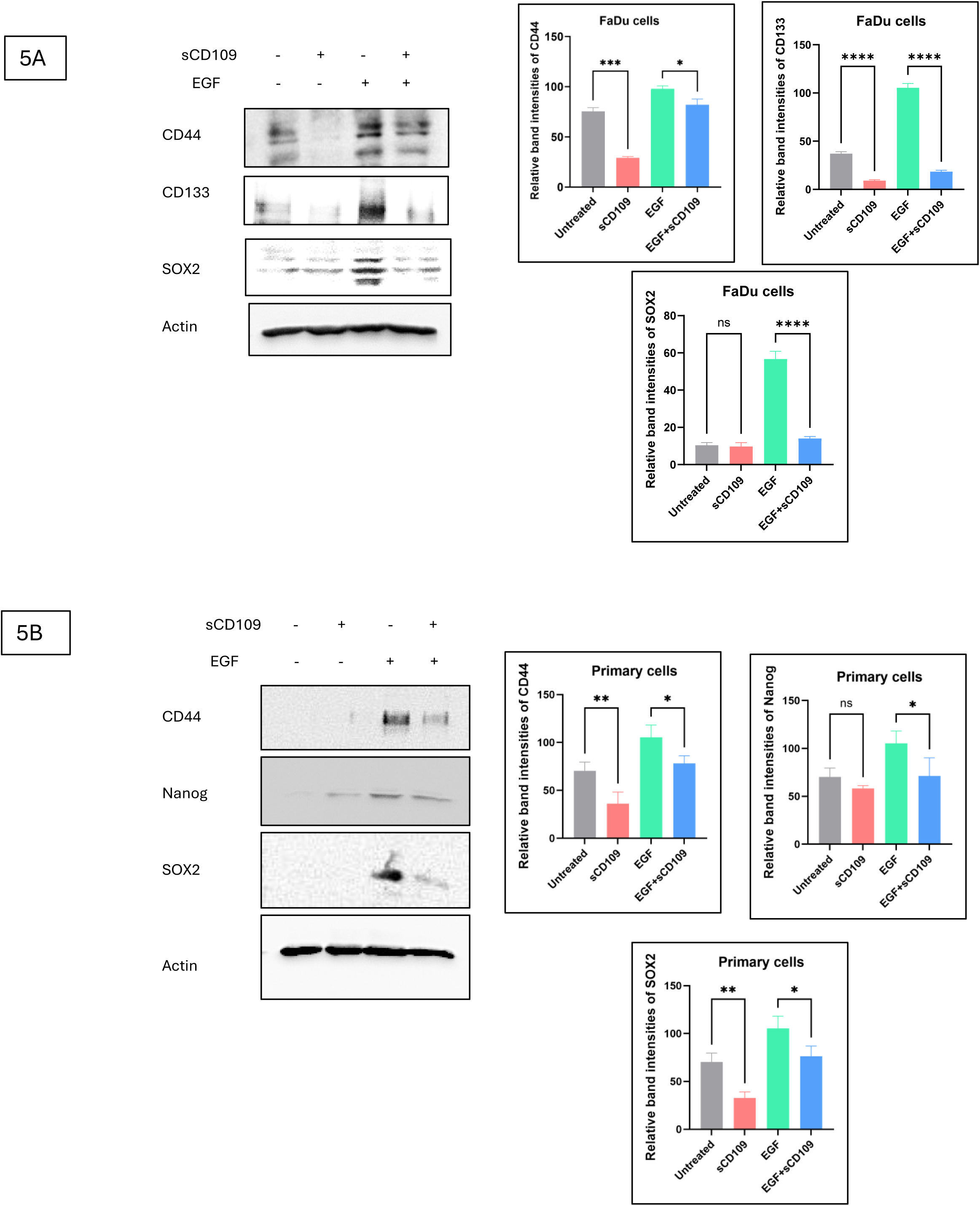
Soluble CD109 suppresses stemness-associated transcriptional programs in HNSCC. A) Western blotting analysis of CD44, CD133, and SOX2 upon treatment of FaDu cells with EGF and/or sCD109 for 48 hours. B) Western blotting analysis of CD44, Nanog, and SOX2 upon treatment of patient 2’s primary cells with EGF and/or sCD109 for 48 hours. All the results were expressed as the mean+/- SD of three independent experiments. Significance was calculated using one-way ANOVA *p < 0.01.

#### Soluble CD109 inhibits EGFR-driven tumorigenic phenotypes, angiogenic potential, and 3D spheroid tumorigenicity in HNSCC cells

To determine whether molecular inhibition of EGFR signaling translates into functional phenotypes, we evaluated multiple oncogenic responses. MTS cell proliferation assays revealed that sCD109 treatment for 24 hours reduced EGF-induced cell proliferation in FaDu cells (Fig. 6A). Circular wound-healing assays revealed significant reduction of migratory capacity of FaDu (48 hours) and UMSCC38 (24 hours) cells (Fig. 6B; Supplementary Fig. 1H). To evaluate tumorigenicity in a three-dimensional context, we performed spheroid formation assays using ultra-low attachment plate conditions. The 9-day sCD109 treatment reduced both spheroid number and diameter (Fig. 6C).

**Figure 6:**
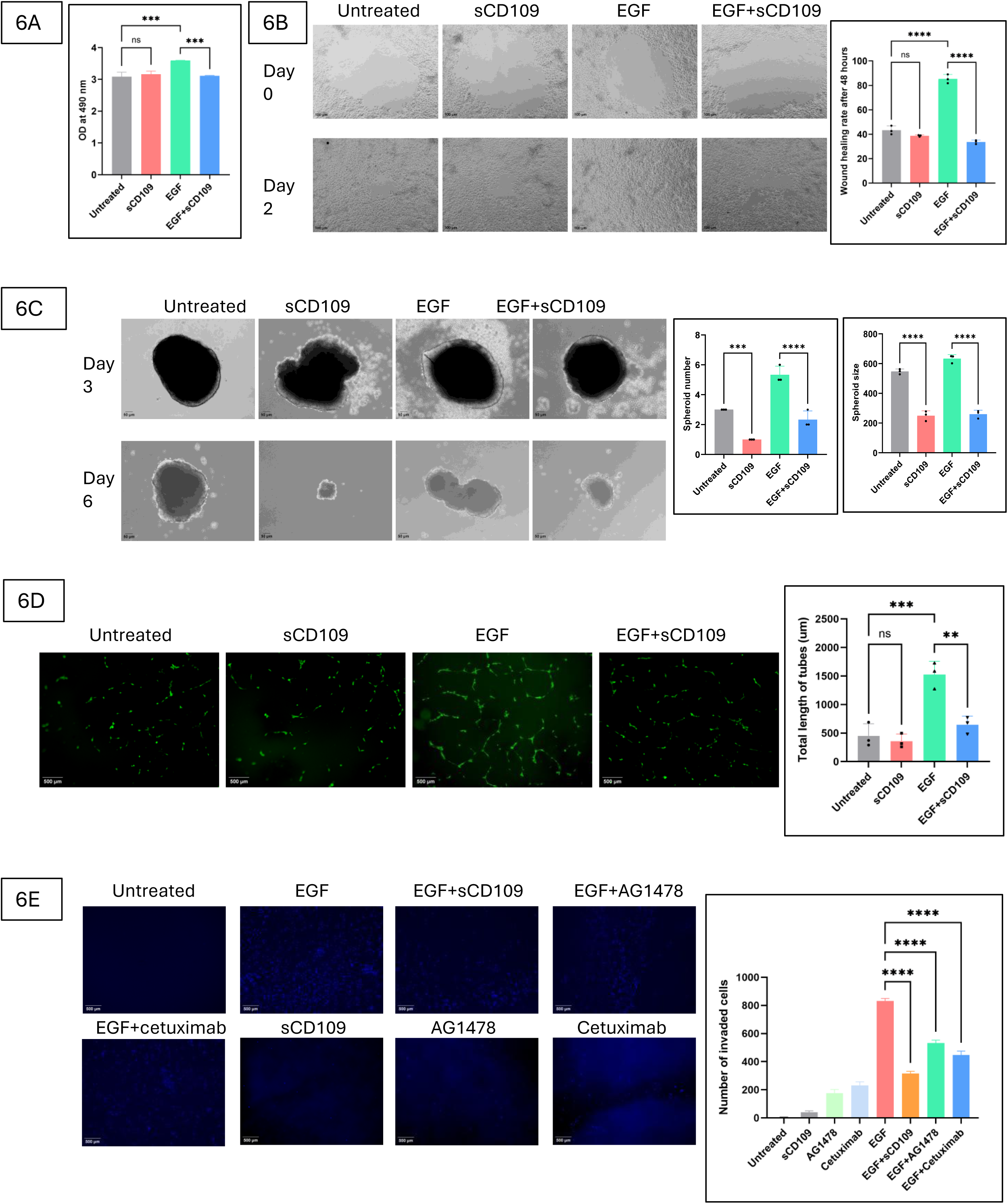
Soluble CD109 inhibits EGFR-driven tumorigenic phenotypes, angiogenic potential, and 3D spheroid tumorigenicity in HNSCC cells. A) MTS cell proliferation assay upon treatment of FaDu cells with EGF and/or sCD109 for 24 hours. B) Circular wound healing assay upon treatment of FaDu cells with EGF and/or sCD109 for 48 hours. C) 3D tumor spheroid formation assay upon treatment of spheroids with EGF and/or sCD109 for nine days treatment time. D) Matrigel cell invasion assay upon treatment of FaDu cells with EGF and/or sCD109, cetuximab, AG1478 for 24 hours. E) Angiogenesis tube formation assay upon treatment of HUVEC cells with EGF and/or sCD109 for 18 hours. All the results were expressed as the mean+/-SD of three independent experiments. Significance was calculated using one-way ANOVA *p < 0.01.

In Matrigel cell invasion assays, sCD109 strikingly reduced the number of invaded cells from top to bottom chambers, compared with EGF-induced controls within 24 hours of treatment (Fig. 6D). The intensity of sCD109’s inhibition on cancer cell invasion was greater than that with pharmacologic EGFR blockade, supporting the functional relevance of sCD109-mediated receptor suppression. Finally, to examine effects on angiogenic potential, we assessed endothelial tube formation in HUVEC cells. The 18-hour sCD109 treatment impaired the formation of capillary-like networks on top of solidified Matrigel, as evidenced by decreased total tube length, when compared to EGF controls (Fig. 6E). Collectively, these functional assays demonstrate that sCD109 exerts anti-tumorigenic potential, suppressing proliferation, migration, invasion, spheroid tumorigenicity, and angiogenic potential in cell models.

### Discussion

The overexpression of EGFR in HNSCC suggests a central role in disease progression (Bhat et al., 2021; Li et al., 2023; Yang et al., 2010). The high rates of locoregional recurrence, distant metastasis, resistance to anti-EGFR therapies and persistently poor survival outcomes in HNSCC patients underscore the urgent need to delineate the molecular mechanisms driving EGFR-mediated tumor progression (Ahmed & Cohen, 2007; Alsahafi et al., 2019; Barham et al., 2025; Shyamsunder et al., 2025).

CD109, a GPI-anchored glycoprotein, is consistently overexpressed across multiple malignancies, including HNSCC, and has emerged as a pro-tumorigenic factor (Cui et al., 2025; Hassan et al., 2025c; Mori et al., 2023; Tsutsumi et al., 2022; Zhou et al., 2022). We recently found that membrane-anchored mCD109 stabilizes EGFR at the cell surface which results in sustained EGFR signaling, upregulating downstream signaling and tumorigenic phenotypes, including enhanced migration, and invasion in HNSCC patient cells (Kungyal et al, 2026/BioRxiv), leading to the abrogation of tumor formation in CD109 KO mice (Zhou et al., 2022).

In contrast to the well-characterized effects of mCD109 on EGFR regulation as mentioned above, the role of endogenously released sCD109 to EGFR signaling has not been defined. Our findings in the current study reveals that sCD109 as a functional antagonist of mCD109’s tumor-promoting activity by suppressing EGFR-driven oncogenic circuitry. Our SPR analysis demonstrates a direct interaction between sCD109 and EGFR and highlights a higher binding affinity of sCD109 for EGFR ectodomain than for homotypic sCD109-sCD109 interaction. Our findings show that, by disrupting mCD109-EGFR interactions, sCD109 reduces cell surface EGFR levels, attenuates activating EGFR phosphorylation at Y1068, enhances phosphorylation at the degradation-associated Y1045 site, and promoting EGFR degradation. Furthermore, sCD109 suppresses EGF-induced nuclear EGFR localization, mCD109-mediated stemness, invasion, and angiogenic potential.

By disrupting mCD109-mediated EGFR stabilization, sCD109 appears to shift EGFR from a signaling-competent state toward receptor downregulation and turnover. Consistent with this model, sCD109 reduces both cell-surface and total EGFR levels in HNSCC cells, as supported by flow cytometry, cell-surface biotinylation with streptavidin pull-down, and immunocytochemistry analyses. This loss of receptor availability accompanies by impaired EGFR activation, as sCD109 attenuates phosphorylation at the signaling-associated Y1068 site, suppresses downstream STAT3, AKT, and ERK signaling, while enhancing phosphorylation at Y1045, a site associated with c-Cbl recruitment, ubiquitination, and receptor trafficking (Capuani et al., 2015). Notably, these effects of sCD109 oppose the actions of mCD109 whose expression was shown to stabilize cell-surface EGFR, promote EGFR phosphorylation at Y1068 and suppress phosphorylation at Y1045 (Kungyal et al, 2026/BioRxiv). Together, these findings suggest that soluble and membrane-anchored CD109 exert opposing effects on EGFR signaling, with sCD109 favoring receptor downregulation and attenuation of EGFR-driven oncogenic signaling.

Beyond limiting EGFR activation at the plasma membrane, sCD109 influences EGFR intracellular fate by restricting nuclear localization and promoting receptor degradation. This is crucial because nuclear EGFRs were implicated in transcriptional regulation, DNA repair, therapeutic resistance, and poor prognosis in several cancers (Li et al., 2010; Liccardi et al., 2011; Zhu et al., 2024). In our study, sCD109 reduced EGF-induced nuclear accumulation of phosphorylated EGFR and accelerated EGFR degradation under ligand-stimulated and unstimulated conditions, respectively. These findings are consistent with literature showing that activated EGFR can undergo multiple trafficking routes, including lysosomal degradation, recycling to the cell surface, or retrograde transport to the nucleus through endosomal, Golgi, and ER-associated compartments (Tomas et al., 2014). Once in the nucleus, EGFR can function as both a co-transcriptional regulator and/or a kinase, targeting PCNA and STAT3 (Brand et al., 2013). In line with cycloheximide chase assays from our group demonstrating that mCD109 overexpression decreases EGFR degradation (Kungyal et al, 2026/BioRxiv), our current data suggest that sCD109 counteracts mCD109-dependent EGFR stabilization and favors receptor turnover. Thus, sCD109 may suppress EGFR oncogenic output not only by inhibiting receptor phosphorylation at the membrane but also by limiting nuclear EGFR localization and promoting receptor degradation.

The inhibitory effect of sCD109 on mCD109-driven EGFR signaling reprograms HNSCC progression. In particular, sCD109 reduces EGF-induced expression of stemness-associated markers, including CD44, CD133, Nanog, and Sox2, contravening our previous findings that mCD109 overexpression promotes EGFR-driven cancer stem-like programs and invasive behaviour (Kungyal et al, 2026/BioRxiv). This is of paramount significance because these markers have been implicated in tumor initiation, plasticity, metastasis, and therapeutic resistance in HNSCC and other cancers (Chen et al., 2018; Li et al., 2017; Liou, 2019). Consistent with this molecular rewiring, sCD109 suppresses oncogenic outputs such as cancer cell proliferation, migration, invasion, 3D spheroid formation, and angiogenic potential, thereby limiting aggressive HNSCC phenotypes.

The biological relevance of sCD109 is supported by our prior studies showing that CD109 can be released from the cell surface through multiple mechanisms, including PI-PLC-mediated release of the GPI-anchored ectodomain (Li et al., 2016; Tam et al., 2001), and furin-mediated processing (Hagiwara et al., 2010). The recombinant sCD109 protein used in the present study, spanning from Valine (22aa) to Serine (1268aa), corresponds to the soluble ectodomain fragment upstream of the furin-processing site, providing a biologically relevant molecule to investigate the role of sCD109 upon EGFR regulation. Elevated soluble CD109 levels in the serum were associated with nodal metastasis and advanced disease in HNSCC patients (Hagiwara et al., 2021) and soluble CD109 levels have been found in the serum of CD109-overexpressing tumor-xenograft bearing mice (Sakakura et al., 2014). These observations support the clinical relevance of soluble CD109 and provide a rationale for studying its function in HNSCC. The mechanisms governing the endogenous release of sCD109 and the relative abundance of membrane-anchored versus soluble CD109 in HNSCC remain to be elucidated.

Together, our findings suggest that CD109 membrane anchorage and soluble fragment release dictate whether it amplifies or restrains EGFR-driven oncogenic programming. This model is reminiscent of other receptor systems in which membrane-bound and soluble isoforms exert distinct biological functions, with soluble receptors modulating signaling through ligand sequestration, trans-signaling, or regulation of receptor availability (Park & Lee, 2024). Such functional divergence has been described for IL-6R and TNF receptor family members and is increasingly recognized as an important determinant of cancer progression and therapeutic response (Park & Lee, 2024).

Collectively, we show for the first time that sCD109 functions as a soluble antagonist of mCD109-mediated EGFR oncogenic signaling in HNSCC, thereby uncovering a previously unrecognized regulatory mechanism within the CD109-EGFR axis (Fig. 7). Through disrupting the stabilization of mCD109-EGFR complexes, impairing EGFR activation, promoting its degradation, and inhibiting stemness-associated and tumorigenic programs, sCD109 acts as a key modulator of EGFR’s oncogenic circuitry. These results provide a strong mechanistic basis for future studies to investigate the regulation of sCD109 release by multiple enzymes, assess the tumor-specific effects of an altered mCD109/sCD109 balance, and examine whether sCD109-mediated rewiring of EGFR oncogenic mechanisms could circumvent EGFR-driven therapeutic resistance in preclinical HNSCC models.

**Figure 7:**
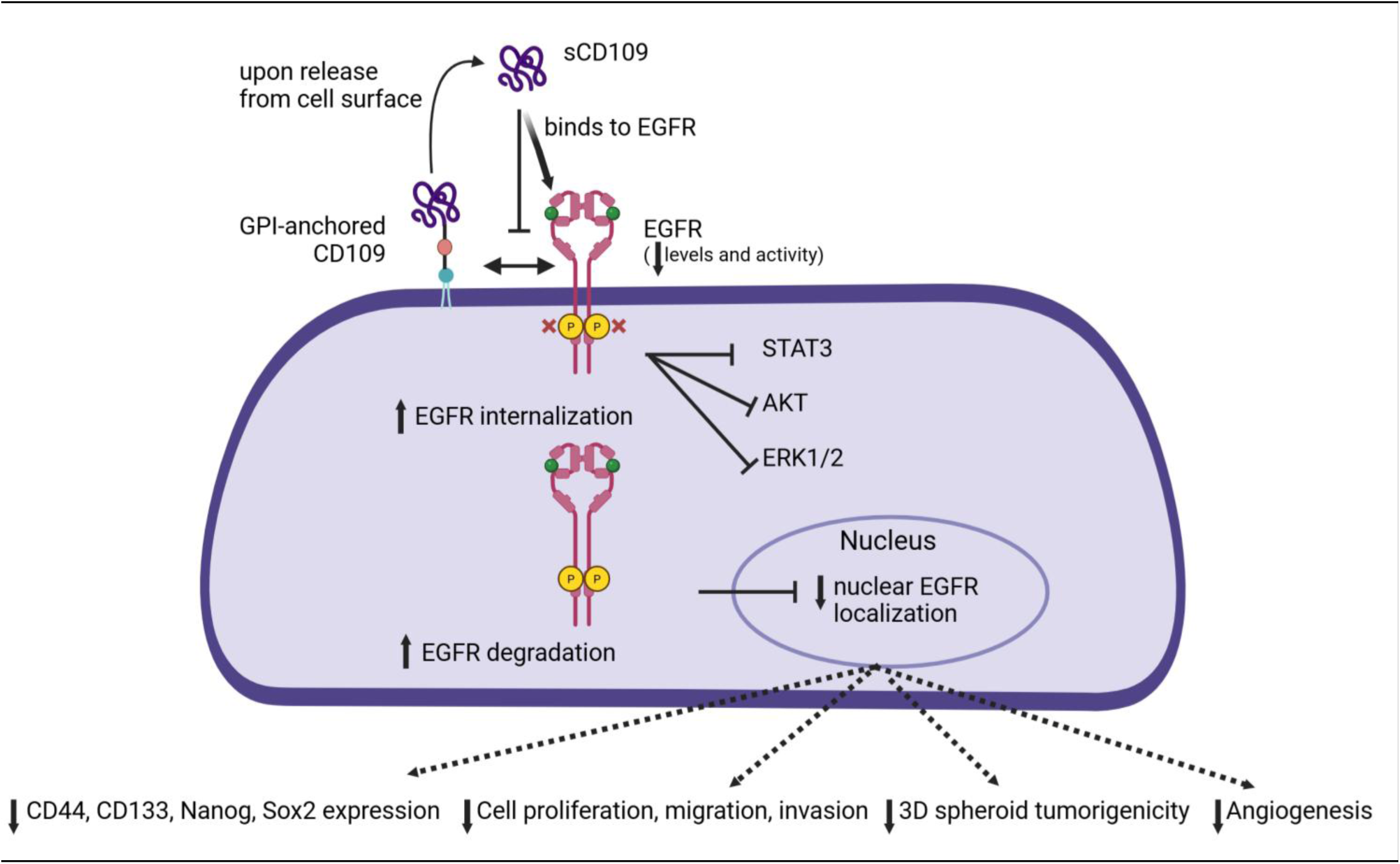
Graphical abstract. Our previous research showed that mCD109 is pro-tumorigenic in SCC. However, our current study shows that sCD109 binds to EGFR with high affinity, thereby inhibiting the interactions of mCD109-EGFR on cell surface. sCD109 reduces the cell surface levels of EGFR thereby inhibiting EGFR phosphorylation at Y1068 site, while promoting phosphorylation at Y1045 site. sCD109 inhibits EGF-induced phosphorylation of STAT3, AKT and ERK1/2 in FaDu cells. sCD109 blocks the nuclear localization of phosphorylated-EGFR while promoting EGFR degradation. Additionally, sCD109 inhibits the EGF-induced expression of cancer stem cell markers – CD44, CD133, Nanog and SOX2. The sCD109 predominantly attenuates the cancer cell proliferation, migration, 3D spheroid tumorigenicity, invasion in FaDu cells and angiogenic potential in HUVEC cells.

## Figure legends

**Supplementary figures. 1A)** Western blotting analysis of phospho-EGFR (Y1068), total EGFR and Actin upon treatment of FaDu cells with EGF for different time points of 0,5,10,15,20,30 mins. **1B)** Western blotting analysis of phospho-EGFR (Y1068), total EGFR and Actin upon treatment of FaDu cells with EGF and/or sCD109 at different concentrations of 1,10,20,50,100nM for 15 mins. **1C)** Western blotting analysis of phospho-EGFR (Y1068), total EGFR and Actin upon treatment of patient 1’s primary cells with EGF and/or sCD109 at different concentrations of 1,5,10nM for 15 mins. **1D)** Western blotting analysis of phospho-EGFR (Y1068), total EGFR, phospho-STAT3 and Actin upon treatment of UMSCC38 cells with EGF and/or sCD109 for 15 mins. **1E)** Circular wound healing assay upon treatment of UMSCC38 cells with EGF and/or sCD109 for 24 hours.

## Authors contributions

Conceptualization: AP, VRD

Data curation: VRD, TK, VN

Formal Analysis: VRD, KF, AP, DPR

Funding acquisition: AP

Methodology: VRD, AH, KF, AP

Project administration: AP

Resources: AP, DPR, NS

Software: VRD, VN

Supervision: AP

Validation: VRD, AP

Visualization: VRD, AP

Writing – original draft: VRD, AP

Writing – review & editing: VRD, KF, DPR, AP

All authors have read and agreed to the published version of the manuscript.

## Acknowledgements

The authors sincerely thank Dr Simon Tran, Dental Medicine and Oral Health Sciences at McGill University for gifting the UMSCC38 cells. We also sincerely thank Dr Chantal Seguin for the gift of HUVEC cells. The authors thank the flow cytometry, Histology and Confocal microscope facilities (Julien Leconte, Min Fu and Shibo Feng) at RI-MUHC and the Goodman Cancer Center, McGill University. The authors also thank Dr Zein Amro for helping with the circular wound healing assay protocol. The study was funded by CIHR project grants, PJT 162132 and PJT 148916 awarded to A. Philip and an SSP/SIS graduate scholarship award and a RI-MUHC studentship award awarded to V R. Durgempudi.

## Data availability statement

No datasets were generated or analysed during the current study.

