## Supplementary data for "Rewiring of EGFR oncogenic program by opposing actions of membrane versus soluble CD109 in HNSCC"

| **Patient's ID** | **Patient 1** | **Patient 2** |
|---|---|---|
| HPV status | p16 unknown | p16 -ve |
| histological diagnosis category | Squamous cell carcinoma | Squamous cell carcinoma |
| Subtype of SCC | Convensional SCC | Convensional SCC |
| anatomical tumor site | Oral Tongue (FOM) | Oral Tongue |
| lateral | Left | Left |
| Sex | Male | Female |
| location | Oral Cavity | Oral Cavity |
| stage | T2 | T2 |
| age | 73 | 58 |
| Diagnosis date | 5-Apr-22 | 27-Sep-22 |
| Chemotherapy treatment | No | No |
| Did patient receive primary radiation | No | No |
| Did patient receive adjuvant radiation | No | No |
| Smoking history CURRENT: Patient has smoked cigarettes within 1 year of their diagnosis PREVIOUS: Patient quit smoking cigarettes at least 1 year prior to diagnosis NEVER: Patient has never smoked cigarettes UKNOWN | Current | Never |
| Pack-years - 1 pack = 20 cigarettes - Eg. 1 pack-year is equal to smoking 1 pack per day for 1 year | 50 | 0 |
| Pack-year stratifying group | >= 10 pack years | 0 |
| Alcohol Never: Never >2 drinks per day for >2 weeks Current: >2 alcoholic drinks per day within the last 2 weeks Previous: Currently drink less than above, but at some point in their lives drinking >2 alcoholic drinks per day for >2 weeks | Previous: | Never |
| Alcohol quantity | ? | 0 |
| Date of surgery | 5-Sep-22 | 11-Oct-22 |
| Type of surgery | L Neck dissection and L hemiglossectomy of L tonsil | L partial glossectomy, L neck dissection |
| surgical path report | pT4a, pN3b | pT3, pN3b |
| Node metastasis | Positive | Positive |
| WPI | Identified | Identified |
| PNI in nodes | Identititifed | Identified |
| LVI in nodes | Identified | Identified |
| ECE in nodes | positive | positive |

### Supplementary figures

1. EGF time curve in FaDu cells
2. sCD109 dose curve in Fadu cells
3. sCD109 dose curve in primary 195 cells
4. pEGFR inhibition in UMSCC38 cells
5. Cell migration in UMSCC38 cells

**Supplementary fig 1**

### EGF time curve for pEGFR (Y1068 and Y1045) in FaDu cells

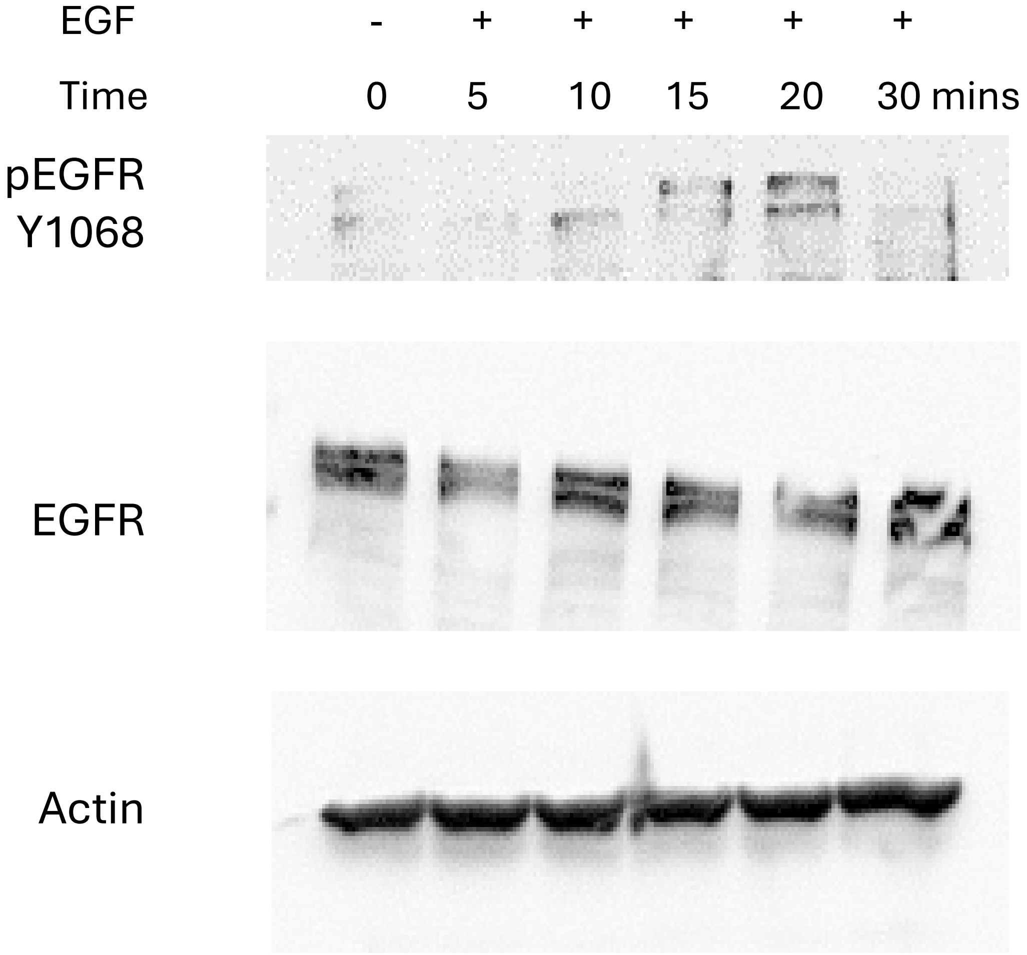

**Supplementary fig 2**

sCD109 inhibits pEGFR in dose-dependent manner in FaDu cells

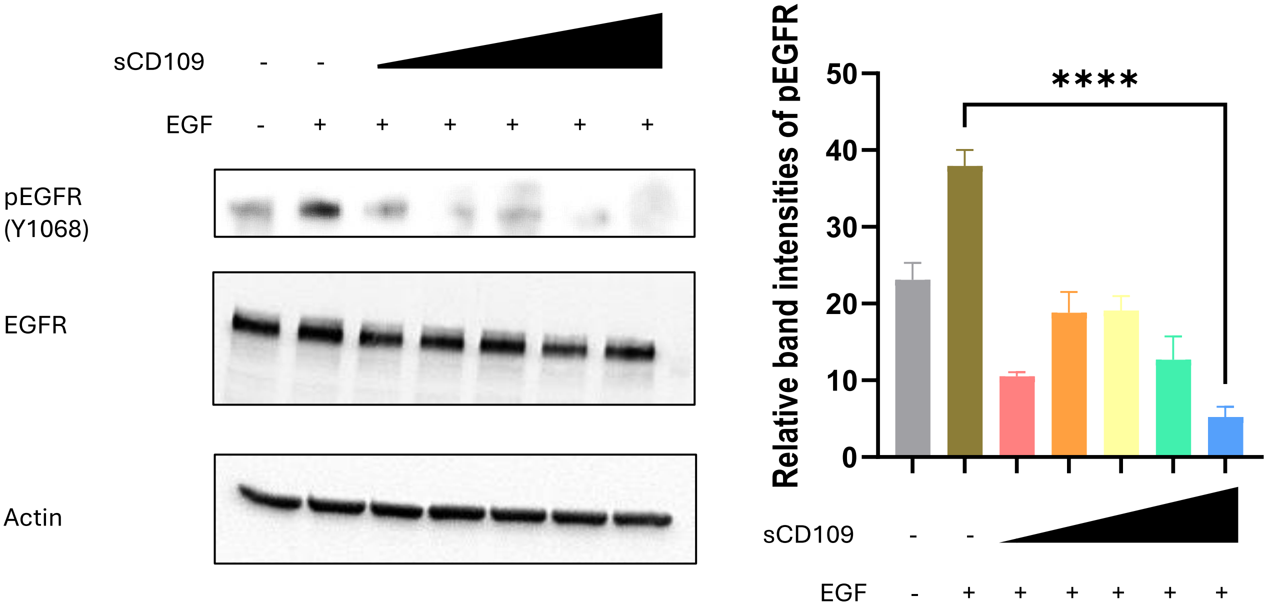

**Supplementary fig 3**

sCD109 dose curve in patient 1's primary cells

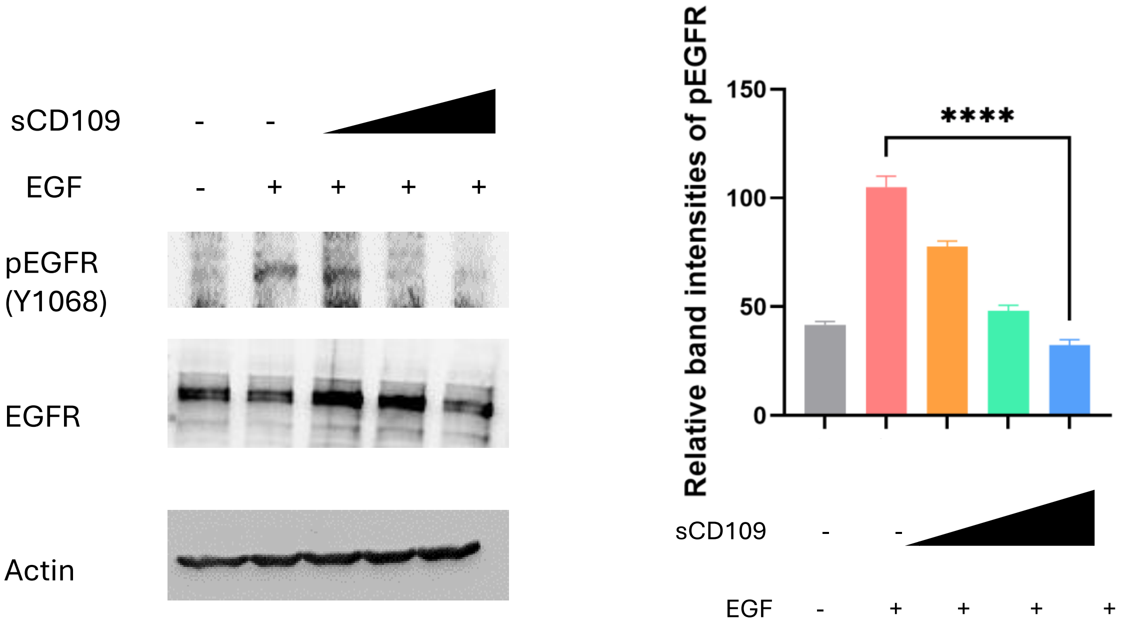

**Supplementary fig 4**

pEGFR inhibition in UMSCC38 cells

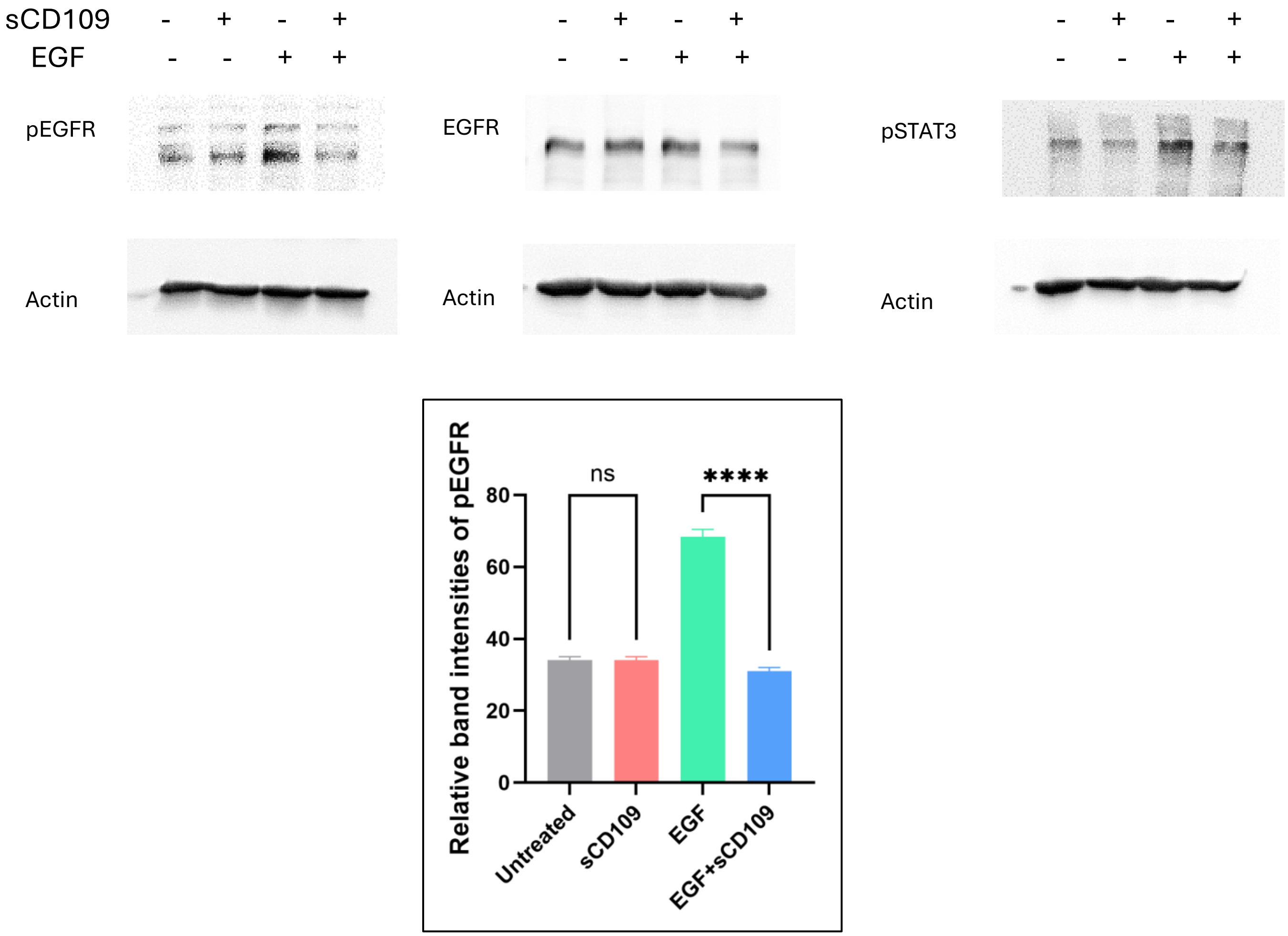

**Supplementary fig 5**

sCD109 inhibits cancer cell migration in UM-SCC-38 cells

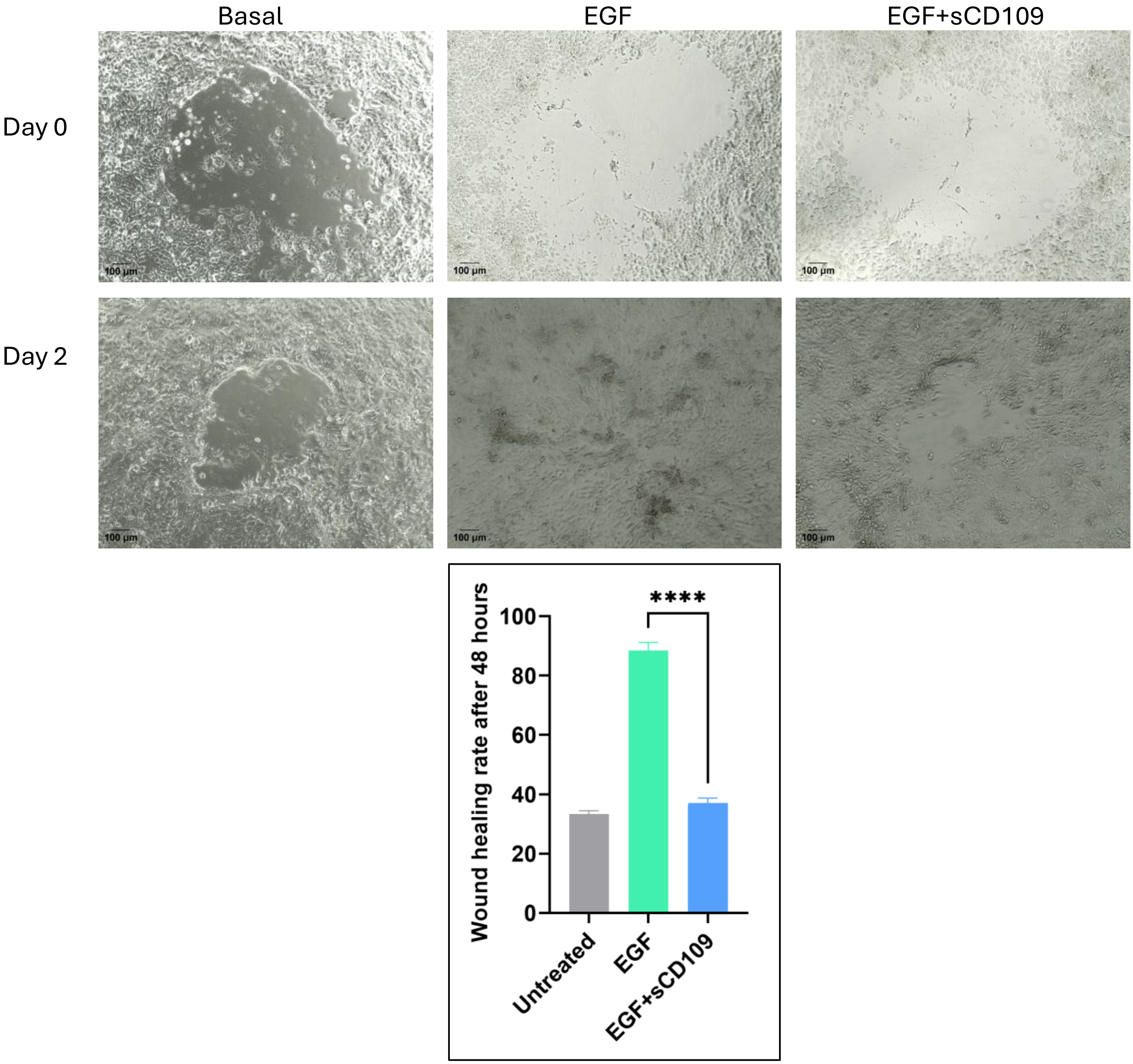
